# G-*PLIP*: Knowledge graph neural network for structure-free protein-ligand bioactivity prediction

**DOI:** 10.1101/2023.09.01.555977

**Authors:** Simon J. Crouzet, Anja Maria Lieberherr, Kenneth Atz, Tobias Nilsson, Lisa Sach-Peltason, Alex T. Müller, Matteo Dal Peraro, Jitao David Zhang

## Abstract

Protein-ligand interaction (PLI) shapes efficacy and safety profiles of small molecule drugs. Existing methods rely on either structural information or resource-intensive computation to predict PLI, making us wonder whether it is possible to perform structure-free PLI prediction with low computational cost. Here we show that a light-weight graph neural network (GNN), trained with quantitative PLIs of a small number of proteins and ligands, is able to predict the strength of unseen PLIs. The model has no direct access to structural information of protein-ligand complexes. Instead, the predictive power is provided by encoding the entire chemical and proteomic space in a single heterogeneous graph, encapsulating primary protein sequence, gene expression, protein-protein interaction network, and structural similarities between ligands. The novel model performs competitively with or better than structure-aware models. Our observations suggest that existing PLI-prediction methods may be further improved by using representation learning techniques that embed biological and chemical knowledge.

## 1 Introduction

Most drug-like small molecules act as ligands binding to proteins [1, 2]. Protein-ligand interaction (PLI) with primary and secondary pharmacological targets critically determine the efficacy and safety profile of a drug [3, 4]. Prediction, quantification, and interpretation of PLI is, therefore, an integrative component of pre-clinical drug discovery [5, 6].

A plethora of experimental methods has been developed for PLI quantification, including well-established biophysical and biochemical binding assays, as well as omics-based methods, such as chemoproteomics [7]. Binding assays can be both fast and accurate for a few targets of interest [8]. However, our ability to probe proteome-wide PLIs with such assays is limited by the availability of large quantities of purified proteins of interest [9]. In addition, quantification of proteome-wide PLIs is labour-intensive, time-consuming, and often results in moderate sensitivity [10].

Computational methods predicting PLIs complement experimental approaches and are indispensable in drug discovery [11–13]. Some methods simulate molecular docking [14, 15]. Other methods rely on structure-based representations of the protein-compound complex, taking solved structures as input for machine-learning algorithms [16–19]. Accumulation of high-quality data derived from biochemical and biophysical screening [20], besides novel algorithms, has empowered such methods, although the underlying limitation, namely only a limited number of complex structures is solved, remains. Nevertheless, screening procedures are progressively being complemented or replaced by modelling procedures [21, 22].

Besides the limitation of structures, most existing computational methods do not consider several biological factors that may contribute to PLI. For instance, most of them treat proteins as isolated entities, ignoring protein-protein interactions that are ubiquitous under physiological conditions [23]. Localization and expression of proteins are usually not considered either, leaving room for new prediction algorithms both able to circumvent existing limitations and able to embed additional knowledge into the prediction process. Such algorithms have the potential to improve the prediction performance and to yield a better understanding of the relative importance of individual factors.

During the last years, applications of deep learning techniques have greatly expanded in drug discovery [24]. In particular, geometric deep learning and graph neural networks (GNNs) have been widely used for molecular representations [25], structure-based drug design [26–29], molecular property prediction [30–33], and protein-ligand binding predictions [34]. Besides its good performance in many tasks, two reasons make GNNs particularly appealing in the context of PLI prediction. First, in contrast to traditional deep learning, geometric deep learning can operate in non-Euclidean spaces, thus able to capture diverse information that is not easily embedded into an Euclidean representation, such as graphs that consist of nodes and edges [35]. Thus, GNNs are able to capture complex and interdependent relationships between biological entities in multiple scales. The benefit is particularly prominent in protein-relevant tasks, such as protein representation [29, 36, 37], protein folding [38], protein design [39, 40], and protein-protein interaction [41–47]. Second, various research groups have developed new techniques to improve the interpretability of GNNs [24, 48, 49]. We also observe that heterogeneous multimodal graphs are well-suited for representing various entities and their interactions within real-world systems, and are therefore an ideal choice for coarse-grained modelling of biochemical processes [50]. Bearing these promises as well as challenges and concerns raised about the use of complex deep learning approaches [16, 51] in mind, we consider heterogeneous GNN a natural candidate for the task of PLI prediction.

Here, we report a novel GNN model termed G-*PLIP* (*Graph for Protein Ligand Interaction Prediction*). G-*PLIP* predicts PLI quantitatively using non-structural features of proteins and ligands. We demonstrate that G-*PLIP* performs comparably with or better than state-of-the-art structure-aware models, while the computational cost remains low thanks to its simple model architecture and to the encapsulation of the chemical sub-space into a single graph. We analyzed the contribution of individual feature types to the model’s performances and demonstrated that primary sequences of proteins and molecular information of ligands contain rich information which can be exploited for PLI prediction.

## 2 Methods

### 2.1 Architecture of a heterogeneous GNN

We constructed a heterogeneous knowledge graph 𝒢 = (*N, E*) featuring two distinct node types *N* : ligands encompass drugs or drug-like molecules *N*_*c*_, and proteins encompass target proteins *N*_*p*_. The graph incorporates three types of links: proteinprotein links embodying physical binary interactions between proteins *E*_*p,p*_, ligand-ligand links reflecting chemical similarity between compounds *E*_*c,c*_, and protein-ligand links representing bioactivities between ligands and protein targets *E*_*p,c*_. *G* is undirected, therefore *E*_*p,c*_ = *E*_*c,p*_. The model adopts a non-linear multiconvolutional neural network with an encoder-decoder architecture: the encoder generates node embeddings *via* convolutional layers, and the decoder predicts bioactivities, as shown in Figure 1.

**Figure 1.**
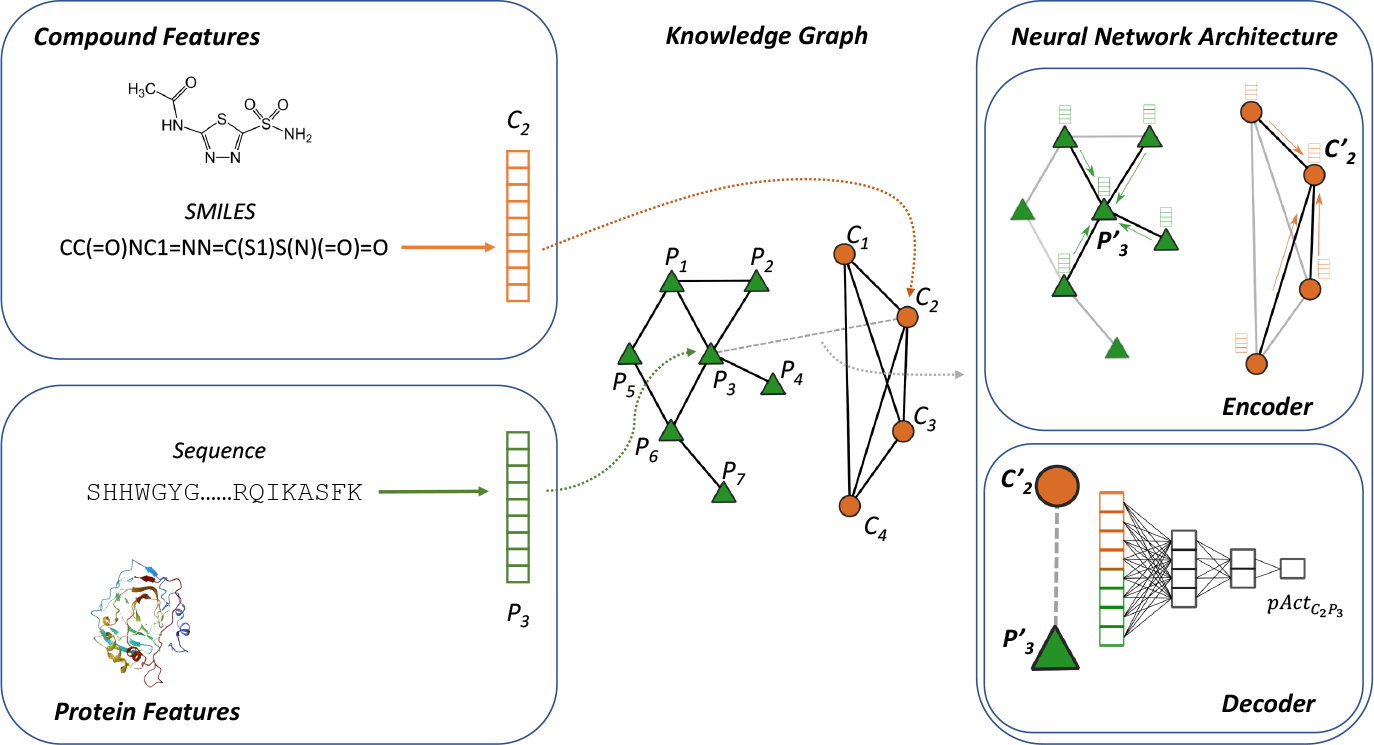
Overview of G-*PLIP*. At its core, the model implements a knowledge graph derived from protein-protein interactome and *in-vitro* pharmacology data (middle panel). The heterogeneous graph consists of binary protein–protein interactions and drug–drug similarities. Information is integrated into the model in the form of feature vectors of protein and molecule nodes (left panels). The highlighted network in the right panel encodes information of node *P*_3_ and *C*_2_ from their neighbours and decodes their embedding to predict the *P*_3_-*C*_2_ bioactivity (*pAct*).

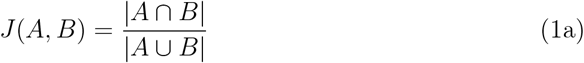

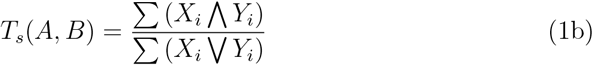

1. The Jaccard and Tanimoto scores.

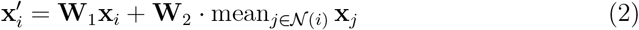
2. The GraphSAGE operator, from Hamilton et al. [52]

#### 2.1.1 Protein features

The encoder leverages three distinctive types of protein features.

- *k*mer count: *k*mer counting from protein sequences empowers G-*PLIP* with an understanding of protein characteristics. This entails tallying the occurrences of every substring of length *k* within a given sequence, yielding a commonly employed distance measure [53]. We hypothesised that, given the relatively modest size of drug-like molecules, it is sufficient to inspect *3*-mers to consider the catalytic site and its immediate vicinity in the context of PLI prediction. Therefore, we used 3-mers as features for a total of 8000 (20^3^) combinations, which were calculated from the protein sequence available on UniProt [54].
- Subcellular location: Proteins’ roles within the cellular machinery can often differ based on their location, thereby influencing their properties and giving insights about their interactions. We used information pertaining to the mature protein’s location and topology in the cell, extracted from UniProt. Subcellular locations are grouped into 31 main classes, and we employ one-hot encoding for representation.
- Gene expression:

Abundance of proteins, or as a proxy, its encoding mRNAs which can be measured by diverse sequencing techniques such as RNA-seq. Gene expression patterns have been correlated with drug effects [55], and can be adapted depending on the cell or tissue type of interest. For each protein, we incorporated normalized Transcripts Per Million (nTPM) from RNA-seq data, averaged across the corresponding expressed genes. We used gene expression data from the Human Protein Atlas [56]. Gene expression was averaged across all tissues by default, although we let this setting open to modification for more specific drug discovery applications to screen compounds interacting with specific pathways.

#### 2.1.2 Compound features

Similarly, the encoder harnesses three distinct types of compound features, each providing a comprehensive grasp of molecular attributes:

- Lipinski’s rule of five: Aiming to evaluate druglikeness, this rule is widely used in pharmacology, assessing whether a compound possesses attributes suitable as an oral drug. The rules [57] state that, in general, an orally active drug has:
  1. Not more than 5 hydrogen bond donors (OH and NH groups)
  2. Not more than 10 hydrogen bond acceptors (notably N and O)
  3. A molecular weight under 500 g/mol
  4. An octanol-water partition coefficient (log P) that does not exceed 5 Rather than to record violations of the rule, which happen increasingly more for certain modern drug classes [58, 59], we chose to input the numerical values of the descriptors proposed above, allowing the model to set its own customized rules form the Rule of Five parameters.
- Chemical features: In order to capture the main characteristics of a compound, we also extracted the following chemical features:
  1. Topological Polar Surface Area (Å^2^)
  2. Number of rotatable bonds (#)
  3. Molecular Refractivity (*MR*) [60]
  4. Number of atoms (#)
  5. Number of aromatic rings (#)
  6. Number of aliphatic rings (#)
  7. Percentage of atoms belonging to aromatic rings (%)
- Fingerprints: We generated Morgan fingerprints [61] from the SMILES representation of compounds and folded them into a bit vector. G-*PLIP* uses by default the extended-connectivity fingerprints (ECFP) with a diameter of 4, *i*.*e*., ECFP4. ECFPs are circular topological fingerprints designed for molecular characterization or similarity searching, being widely employed as molecular descriptors for machine learning models [11]. Those identifiers represent substructures present within the molecule, by assigning integer identifiers to each non-hydrogen atom of the input molecule and iteratively updating them to capture larger and larger circular neighbourhoods. Those final integer identifiers can be converted to fixed-length bit strings, with the risk of collision and therefore loss of information increased as the bit string is shorter. As neural networks can be quite sensitive to background noise, we encoded our fingerprints in a 256 bits vector [62, 63]. The size and the choice between ECFP and FCFP (Functional-Class Fingerprints [61]) of the model can be configured by the user.

With the default setting, the graph encapsulates 8032 features for the protein nodes *N*_*p*_ and 267 features for the compound nodes *N*_*c*_.

#### 2.1.3 Interaction networks

We extracted 15,749 protein-protein interactions (PPIs) between 939 human proteins from a PPI database successfully used by [50] on a similar heterogeneous graph neural network approach investigating polypharmaceutical side effects. We linked all the ligands together to form a full graph, where ligand-ligand connections symbolize compound similarities quantified by Tanimoto scores (Equation 1b), a measure resembling the Jaccard Index (Equation 1a). The Jaccard index measures similarity between finite sample sets and is defined as the size of the intersection divided by the size of the union of the sample sets. Likewise, the Tanimoto score (also referred as Tanimoto similarity ratio) expresses a similarity ratio over Morgan fingerprint bitmaps, defined as the number of common bits divided by the bit length. Similarity values range from 0 to 1, where 1 indicates identical molecules (accounting for potential information loss during bit encoding). Our pipeline permits customization of ligand connectivity to consider only connections surpassing a specified Tanimoto score threshold. By default, we incorporated all compound-compound similarities (keeping the threshold to 0.0), giving a complete subgraph of compounds. It results in a graph with *C* + *P* nodes and 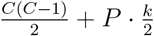 edges, with *P* representing the number of proteins, *C* the number of ligands and *k* the average degree of connectivity of proteins, defined as 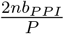 (since the graph is undirected) depending on the PPI database used. In the default configuration, the graph has two distinct connected components, thus we attempted to model the protein-ligand edges in one of the experiments (Table 6, reported as ‘w/ DTIs’).

### 2.2 Link predictions

The proposed GNN consists of two primary components: (a) an encoder producing node embeddings from the original graph, and (b) a decoder using embedding features to forecast drug-target activities. The encoder processes an input graph *𝒢* and generates a *d*-dimensional embedding for each node, aiming to incorporate original node information. This encoder contains two distinct components: the first encapsulates proteins and binary protein interactions, while the second handles ligands and compound-compound similarities.

Both components of the encoder consist of a singular convolutional layer, updating the node features within a latent dimension using the GraphSAGE operator (Equation 2) designed for low-dimensional embeddings of nodes in large graphs [52]. The GraphSAGE operator serves as learnable function that generates embeddings by aggregating information from a node’s local neighbourhood. As message-passing GNNs cannot be applied trivially to heterogeneous graphs, we bypass this by having individual convolutional layers for each edge type. Therefore, the algorithm iterates over edge types to perform the convolution.

The decoder is composed of two linear layers, which employ concatenated embeddings of the protein target and compound molecule to predict bioactivity as a continuous value. A dropout layer is interposed between the two linear layers to prevent overfitting. Given that bioactivities are commonly expressed in logarithmic scale (pAct, defined as -log10(Ka)), we used the Rectified Linear Unit (ReLU) activation function to rectify any negative values within the layers that wouldn’t align with biological meaning.

#### 2.2.1 Model training

We formulate the task of PLI prediction as a multirelational link prediction problem in a heterogeneous graph. We subsequently trained a GNN model to predict bioactivities of unseen protein-ligand pairs (Figure 1), using the root mean square error (RMSE) as the loss function for model training and backward propagation. We opted for a regression task, primarily due to the inherent subjectivity associated with any threshold value used to categorize ground truth data, and secondly because of the goal to emphasize possible off-target effects by directly comparing bioactivities. The model possesses 8449 trainable parameters, and was trained using the Adam optimizer [64] with a learning rate of 0.001 and a batch size of 512 over 500 epochs. Diverse configurations have been assessed during the design process, notably different numbers of convolution layers, types of convolution layers, learning rates, batch sizes, dropouts, PPI networks, compound similarity thresholds, or sizes of Morgan fingerprints (Table S3, S1a and S1b).

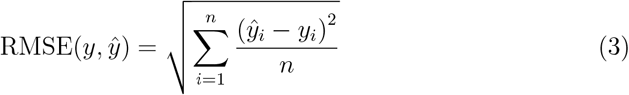

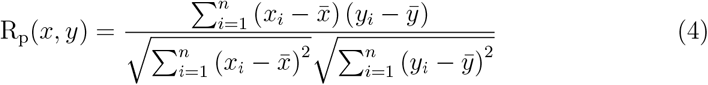

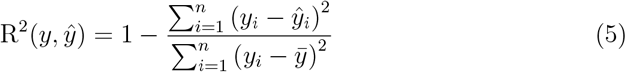

RMSE stands as the prevalent loss metric for regression models and is conventionally used to assess model performances, whereas we also reported several other metrics. Commonly used, the Pearson’s correlation coefficient (*R*_*p*_) gauges the linear correlation between the ground truths and the predicted values. Nonetheless, its exclusive focus on correlation can yield diverse interpretations and has been a subject of concern in regression evaluation [65]. Consequently, we also reported the coefficient of determination *R*^2^.

Furthermore, for accurate identification of off-target activities, the ranking of predicted bioactivities in relation to one another is sometimes as important as exact predicted values, because the rankings allow prioritization of off-target testing and generation of hypotheses for drug repurposing. In essence, identifying off-target effects provides new insight into existing drugs, unveiling their potential to treat diseases for which they were not initially designed, for instance in the case of the repurposing of the drug rapamycin [66]. Therefore, we used a new metric, Rank-Biased Overlap (RBO) [67], to evaluate G-*PLIP*’s proficiency to prioritise likely off-target effects. Compared to Spearman’s *ρ* commonly used to evaluate rankings, RBO presents the advantage of being weighted and favours the top of the ranking depending on its persistence parameter *p*, set as 0.5 for this study.

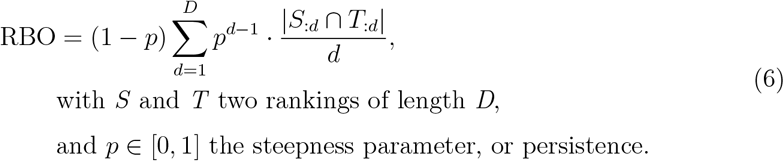

Designed to measure similarity between nonconjoint, indefinite or incomplete ranking lists, RBO is tuneably top-weighted (parameter *p*) and provides a consistent way to handle tied ranks and prefix rankings of varying lengths.

### 2.3 Datasets

#### 2.3.1 The Roche PACE dataset

We trained the model on data derived from the Roche Pathway-Annotated Chemical Ensemble (PACE) library, an extension of the published Roche Small-molecule PAthway Research Kit (SPARK), a chemogenomic library annotated with biological pathways[68]. We leveraged the data that we collected for the PACE library to train G-*PLIP*, leveraging 2972 proprietary PLI measures for 2057 PACE compounds.

Since it is easier to predict the affinity of compounds that are derivatives of a well-studied pharmacophore than to make predictions for new compounds, we also performed scaffold splitting using the Bemis & Murcko (BM) scaffolds [69]. As scaffolds serve as the foundation for the subsequent exploration of derivative compounds designed and synthesized based on their structural framework, scaffold split gives a more realistic estimate of a model’s performance for drug-discovery purposes. We divide compounds into R-groups, linkers, and ring systems, and define a scaffold by removing all R-groups while retaining all ring systems and linkers between them. We use scaffolds to identify molecules that belong to the same chemotype and split into train and test sets that do not overlap in these terms. This approach ensures maximal dissimilarity of train and test sets and reflects a realistic drug discovery scenario where models are applied on previously unseen scaffolds.

#### 2.3.2 The PDBBind dataset

The PDBBind dataset, curated by Wang *et al*. [70, 71], presents a comprehensive collection of qualitative binding affinity data for biomolecular complexes, enriched with detailed structural information. PDBBind serves as one of the gold standards used to assess structure-based PLI models [16–19], such as for the popular Comparative Assessment of Scoring Functions (CASF) benchmark. The general set of PDBBind 2019 includes 17652 protein-ligand complexes, with 4852 of them being in the refined set which contains complexes with high-quality structural data. Within this database, the PDBBind Core set aims at providing a relatively small set of highquality protein-ligand complexes for evaluation and comparison of docking/scoring methods and is notably used as the final evaluation set for the CASF competition. The previous version of this core set, used for CASF-2016, contains 285 protein-ligands complexes from 57 target proteins.

We evaluated the overlap between the PACE dataset and the PDBBind dataset. We observed that the two datasets are highly divergent, with a Jaccard index (Equation 1a) of 0.02 between available compounds (207 compounds) and of 0.004 between known compound-target bioactivities (41 samples).

### 2.4 Model inference

Subsequently, we scrutinized the informativeness of features of by G-*PLIP*. We examined the relative contributions of individual feature types for prediction, by iteratively comparing multiple instances of G-*PLIP* and investigated the impact of different features. Given the vast number of possible feature combinations, we restricted our experimentation on (a) informativeness of features in a graph; (b) informativeness of subgraph connectivity with built-in features; (c) informativeness of subgraph connectivity without any features; (d) usage of known drug-target interactions within the graph itself; (e) usage of different interactomes; (f) usage of different gene expressions; (g) exclusive focus on proteins or on compounds; (h) relevance of a simplest representation. We either started from a null model that uses the correct topology but does not contain any informative features and subsequently added features (Table 5 and 6b), and started from the complete model from where we performed an ablation study by removing some of its components (Table 6a).

We reported the four metrics (RMSE, RBO, *R*_*p*_, and *R*^2^) alongside the improvement, defined by the formula 7 expressing the improvement from the null to the complete model.

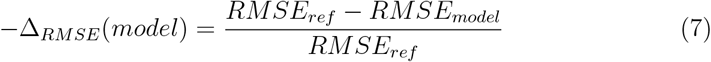

To distinguish the learned biochemical insights of the model from any artificial correlations learned from graph topology, we defined a null model as trained on randomly labelled edges, with bioactivities drawn from a uniform distribution between 3 and 11 (pAct values considered as extremes). To quantify stochasticity, we repeated the experiment 20 times with different random seeds and reported the averaged results as being representative of the null model.

We then tested G-*PLIP* on a dataset of randomly connected edges, with bioactivities shuffled alongside the known drug-target pairs from the existing data, and performed the experiment with the seed used for further tests relative to feature informativeness (Table 1). This procedure aimed to distinguish potential bias stemming from the dataset curation protocol from the model’s actual learning ability.

**Table 1:**
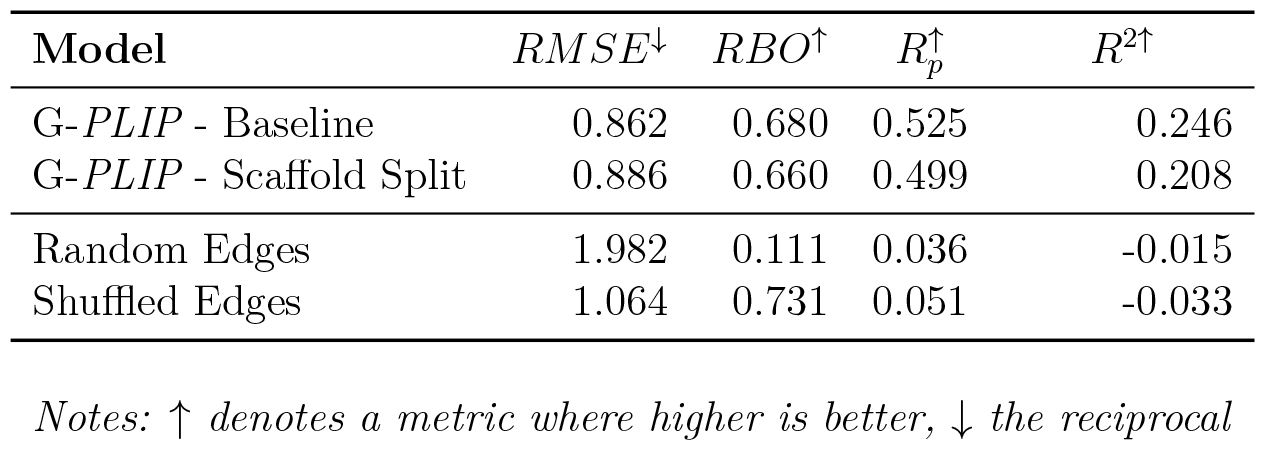
G-*PLIP* assessment on the Roche PACE dataset using different settings, depicting the difference of performances between the default stratified split, the scaffold split, and two control conditions (edge labels randomly assigned, and edge labels shuffled).

Finally, we investigated concerns raised by previous studies about the relevance of intricate deep learning methodologies compared to standard techniques for inferring binding affinities [16, 51, 72–74] by comparing the performance of G-*PLIP* to state-of-the-art machine learning models. We thus used AutoGluon, a framework encapsulating state-of-the-art techniques and automating their optimization [75], inputting the same features to evaluate the gain of a graph neural network architecture.

## 3 Results

### 3.1 Prediction of drug-protein activity (pAct)

G-*PLIP* was primarily trained and evaluated on data derived from the Roche Pathway-Annotated Chemical Ensemble (PACE) library. We first performed a 80%-20% split, stratifying the split by proteins. Each of the proteins will have the same proportion of links dispatched between the train and test set, for an equal representation across the sets to guarantee a fair evaluation. Proteins possessing only one link were excluded from the split and retained for a specific use case. We then used a 80%-20% scaffold split to evaluate the model, grouping molecules derived from the same scaffold into the same set. G-*PLIP* produced a good accuracy in both settings (Table 1), particularly noteworthy with respect to its performances when subjected to randomized data.

We investigated the model’s behaviour and observed intriguing characteristics. First, despite largely concordant predictions, the model displays a propensity to underestimate high bioactivities on the Roche PACE dataset (Figure 2, panel a), without being subsequently observed in the PDBBind dataset (Figure 2, panel b and c). The tendency leads to discrepancy between the performance judged by different metrics, with *R*_*p*_ and *R*^2^ scores indicating a moderate performance while *RMSE* indicating a good performance. Second, the model is also able to predict *pAct* values for missing links, i.e. cases where there is no quantitative information about the interaction between a protein and a ligand. Note that in the context of protein-ligand interaction, a missing link does not necessarily mean a lack of interaction; it may arise due to a lack of measurement, a reporting bias, due to the sensitivity of the assay, or a combination of these factors. Interestingly, the median predicted value of missing links is lower than the predicted values for non-missing links (Supplementary Figure S2). Finally, our model achieves a comparable performance for a wide range of targets classified by their molecular functions (Figure 3 and Supplementary Figure S1).

**Figure 2.**
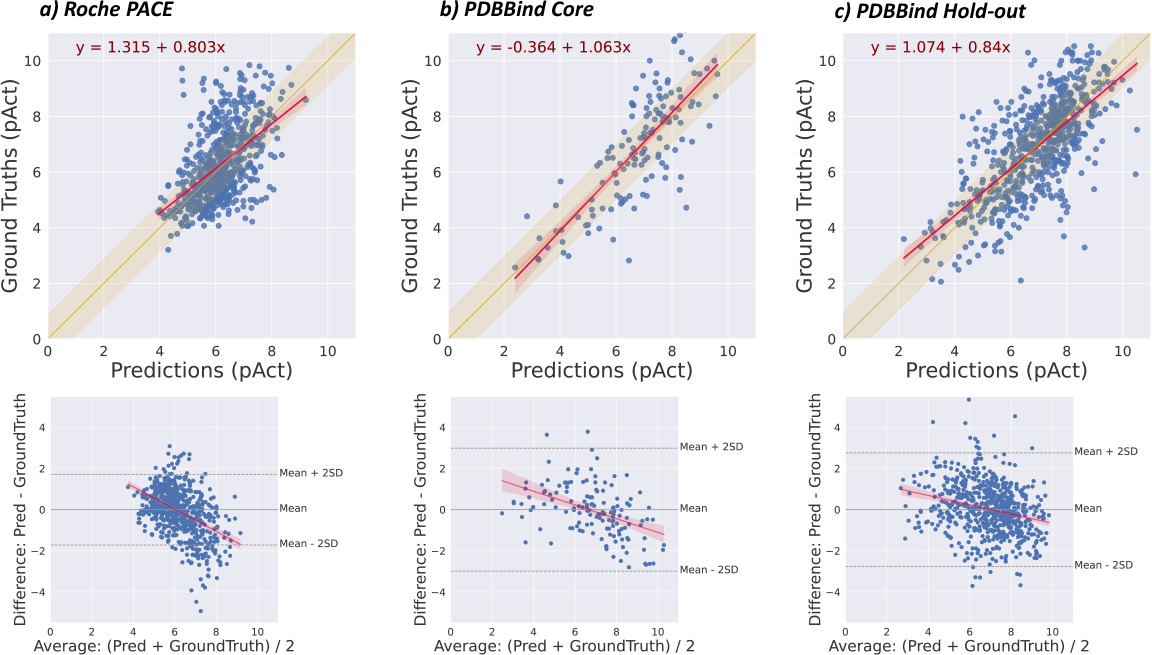
Performance of G-*PLIP* with PACE and PDBBind datasets. Top panels are scatter plots of ground-truth values (*y*-axis) against predictions (*x*-axis) from the test samples. In yellow is highlighted the area of values below the mean *RMSE*. A fitted linear regression model prognosticating ground truths from predictions is sketched in red, with 95% confidence intervals indicated in shades. In the bottom panels, MA plot portraying the differences between predicted values and ground truths (*y*-axis) against the average of the two (*x*-axis). Mean and two units of standard deviation of the differences are indicated in dashed lines. Trendline to visualize the bias is highlighted in red. Displayed are test samples from the Roche PACE dataset (panel a), the PDBBind 2016 CASF Core set (panel b) and the PDBBind 2019 Hold-out set (panel c).

**Figure 3.**
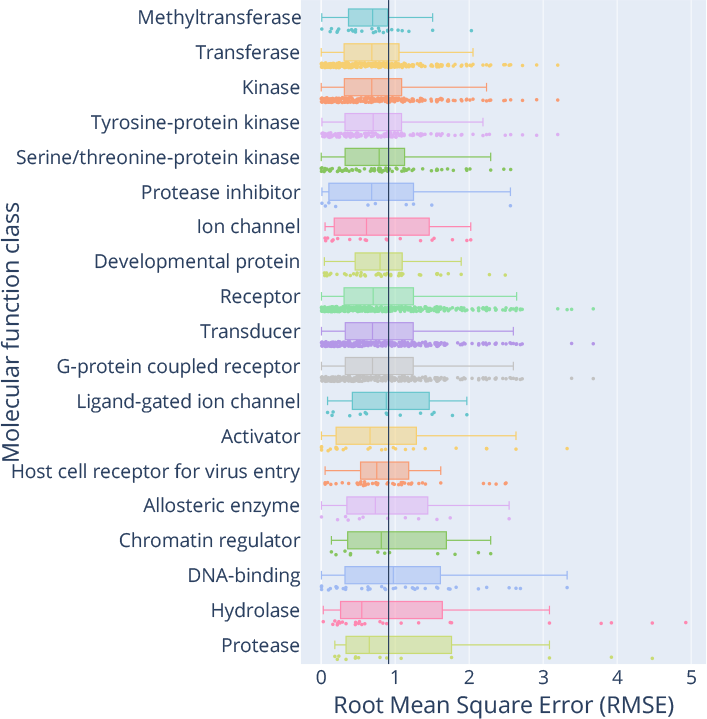
G-*PLIP*’s performance, trained and tested with the PACE database, stratified by molecular function classes of targets. Target classes are sorted by median *RMSE* values. The black vertical line is the overall mean value across all molecular function classes. Molecular functions possessing less than 10 targets are not displayed.

### 3.2 Comparison with state-of-the-art pipelines

Having established a baseline performance, we next evaluated G-*PLIP* with independent datasets and compared it with other pipelines designed for the same task on *PDBBind*, a collection of independent datasets curated by Wang *et al*.[70, 71]. We compared the performance of G-*PLIP* with those of cutting-edge models that leverage structural information [16–19], primarily by using convolutional neural networks or recurrent neural network-based approaches. In order to assess the merit of the structure-free methodology, our model only leveraged the amino-acid sequence of proteins and the SMILES representations of compounds despite the availability of structural data in the *PDBBind* dataset. Since our approach is grounded in human tissue gene expression data and tailored for applications in drug discovery, we confined the comparison to PLI of human proteins. Employing protocols established by prior studies [16–19], we ran a comparative study with two distinct setups and reported RMSE and R_p_, as they are the metrics reported in the prementioned research works.

First, we assessed G-*PLIP* using as a test set the 2016 CASF Core Set (n=141), a benchmark set compiled by the *PDBBind* team, with the remaining database entries constituting the training set (n=8921). Subsequently, we evaluated G-*PLIP* by a temporal splitting of the *PDBBind* dataset (denoted as the 2019 Hold-out Set), where the 2016 version serves as a training set, while the samples introduced in the 2019 and 2020 versions were utilized to test the model (n=667). We furthermore partitioned our training data into both a training and validation subset, with the latter aiding in the selection of the most optimal model across hyperparameters. Yuel [76] was retrained and assessed using the same train/test split, while K_*DEEP*_ [19], Fusion [18], Pafnucy [17] and MPNN-PLI [16] were obtained from the original publications.

G-*PLIP* outperformed its peers on the external 2016 CASF Core Set judging by RMSE, as shown in Table 2, with an improvement of 23.2% compared to Yuel [76], 19.3% compared to K_*DEEP*_ [19], 21.7% compared to Fusion [18], 28.3% compared to Pafnucy [17] and 32.2% compared to the complete MPNN model (with merged protein, ligand and interaction graph, namely MPNN-PLI) [16]. On the 2019 Hold-out set, G-*PLIP* performs better by 31.3% compared to K_*DEEP*_, 26.8% compared to Fusion, 40.4% compared to Pafnucy [17] and 33.9% compared to the complete MPNN model, yet 5.5% worse than Yuel. In both cases, we did not observe the tendency of underestimating high-affnity interactions as we observed in the PACE dataset (Figure 2, panel b and c). To ensure the consistency of the model’s performance, we conducted three distinct training runs and also reported the standard deviation (SD), confirming its reliability.

**Table 2:** Performances of G-*PLIP* on publicly available datasets over three independent trainings (reported as *mean ± sd*), and comparison with state-of-the-art models. Results for Yuel [76] were manually compiled from a model retrained locally using the parameters proposed by the authors on the same train/test splits. Results for K_*DEEP*_ are reported according to the performances claimed in the original paper [19] (for the 2016 Core Set) and manually investigated by a later study [18] (for the 2019 Hold-out Set). Results for Fusion are reported according to the best performances reached during the study, namely by the *midlevel fusion* [18]. Performances for Pafnucy [17] and for MPNN-PLI were obtained from the original MPNN-PLI paper [16] Comparison on PDB Bind.

| Comparison on PDB Bind |  |  |  |  |
| --- | --- | --- | --- | --- |
| Model | 2016 CASF Core Set |  | 2019 Hold-out Set |  |
| | $RMSE^{\downarrow}$ | $R_p^{\uparrow}$ | $RMSE^{\downarrow}$ | $R_p^{\uparrow}$ |
| *G-PLIP* | **1.024 ± 0.013** | 0.781 ± 0.003 | 0.979 ± 0.011 | 0.679 ± 0.008 |
| Yuel | 1.334 | 0.782 | **0.928** | **0.857** |
| MPNN-PLI | 1.511 | 0.813 | 1.481 | 0.652 |
| Pafnucy | 1.429 | 0.773 | 1.642 | 0.456 |
| $K_{DEEP}$ | 1.270 | **0.820** | 1.424 | 0.487 |
| Fusion | 1.308 | 0.810 | 1.338 | 0.545 |

Pearson correlation coefficients are reported to be slightly lower for G-*PLIP* on the core set, however it has been shown this metric can be illusory to evaluate predictive performances on regression tasks [65], is highly sensitive to extreme outliers and influenced by the range of observations [77]. In consequence, satisfying Pearson correlation coefficients could be reached with models unable to predict relevant values, and this hypothesis was corroborated by training a model using the aforementioned correlation coefficient as a loss function. It yielded a Pearson correlation coefficient of 0.79 while proposing a RMSE of 6.12, and thus does not offer any practical predictive power (Supplementary Figure S4). We also attempted transfer learning by training a model on the PACE dataset and evaluating it on the PDBBind Core and Hold-out test sets. The performance, however, was significantly worse than from the model trained on the PDBBind dataset.

### 3.3 Contribution of low-quality data to improve G-*PLIP* ‘s learning abilities

Previous studies remained inconclusive about the question of whether extending datasets by integrating low-quality affinity data would enhance the performance of machine-learning models. While some studies demonstrated improvements in this context [78], recent investigations have posited that such integration exerts minimal [19] or even misleading influence on model outcomes [73, 79, 80]. *PDBBind*’s general dataset is proposed with a subset curated by the authors, the *refined* set, containing data considered as being especially qualitative. To address this discrepancy, we conducted experiments using distinct subsets from the *PDBBind* collection: the *complete, refined*, and *other* sets. The *complete* set represents the entire *PDBBind* dataset, the *refined* set encompasses a subset of high-quality data curated by the authors, and the *other* set comprises the data not present in the *refined* subset.

Each of those three sets was employed for training and subsequently evaluated on both the 2016 CASF Core Set and the 2019 Hold-Out Set, following the same protocol as previously described. The first setup provides a consistent benchmark using reliable test samples carefully selected and segregated from the three subsets (141 bioactivity values). In the second setup, we aimed to gauge the model’s ability to generalize within the given subset. Here, the test data’s composition directly mirrors the training subset’s characteristics, enabling us to observe the model’s behaviour when trained and evaluated on data of similar quality.

Results indicate that augmenting the dataset with lower-quality samples confers little benefit to the model’s performance (Table 3), and the model appears to capitalize only modestly on the enlarged dataset size. Indeed, when evaluating the model on an external reference set (CASF-2016 Core), the results achieved on the *complete* set are only slightly better than those on the *refined* subset. Additionally, the model trained on the *other* subset fails to reach their baseline performances, despite demonstrating favourable metrics on the hold-out set. The model’s performances displayed from the temporal split appear to be influenced by other factors, possibly including the narrower distribution of bioactivity values. In conclusion, our experiments offers little evidence that including low-quality data improves G-*PLIP* ‘s learning abilities

**Table 3:** Performances of G-*PLIP* on the full set and two subsets of the PDB Bind dataset over three independent trainings (reported as *mean ± sd*). The 2016 CASF Core Set is external and thus constant over the subsets, while the 2019 Holdout Set is done by temporal splitting over the selected subset and thus dependent of its content.

| Model | 2016 Core Set |  | 2019 Hold-out Set |  |
| --- | --- | --- | --- | --- |
| | $RMSE^{\downarrow}$ | $R_p^{\uparrow}$ | $RMSE^{\downarrow}$ | $R_p^{\uparrow}$ |
| Complete | $1.024 \pm 0.013$ | $0.781 \pm 0.003$ | $0.979 \pm 0.011$ | $0.679 \pm 0.008$ |
| Refined | $1.138 \pm 0.020$ | $0.731 \pm 0.005$ | $1.089 \pm 0.021$ | $0.754 \pm 0.005$ |
| Other | $1.495 \pm 0.006$ | $0.659 \pm 0.000$ | $1.003 \pm 0.022$ | $0.590 \pm 0.013$ |

**Table 4:** G-*PLIP*’s performances compared over possible variants.

| Models | $RMSE$ | $RBO$ | $R_p$ | $R^2$ |
| --- | --- | --- | --- | --- |
| Compounds-focused | 0.956 | 0.661 | 0.377 | 0.122 |
| Proteins-focused | 0.914 | 0.598 | 0.444 | 0.161 |
| only $kmer$ & Fingerprints | 0.898 | 0.672 | 0.493 | 0.172 |
| Complete | 0.858 | 0.676 | 0.525 | 0.251 |

**Table 5:** G-*PLIP*’s performances on PACE dataset with different sets of features activated. ‘Null’ uses the graph topology but neither protein nor compound features. Indicated are the sets of features used in addition to the null model. Protein features include *k*mer counts (K), gene expression (GE), and subcellular localization (SL). Compound features include fingerprints (F, for instance extended connectivity fingerprint, ECFP), Lipinski features (L) and other chemical features (Ch). *−*Δ_*RMSE*_ represents the improvement of performance compared with the null model.

| Model | $RMSE$ | $-\Delta_{RMSE}$ | $RBO$ | $R_p$ | $R^2$ |
| --- | --- | --- | --- | --- | --- |
| Null | 1.065 | (*ref*) | 0.596 | 0.061 | -0.034 |
| (K) | 0.915 | +14.1% | 0.596 | 0.438 | 0.141 |
| (GE) | 0.948 | +11.0% | 0.595 | 0.398 | 0.125 |
| (SL) | 0.991 | +6.9% | 0.595 | 0.309 | 0.062 |
| (K + GE) | 0.911 | +14.5% | 0.596 | 0.448 | 0.151 |
| (K + SL) | 0.914 | +14.2% | 0.596 | 0.444 | 0.157 |
| (GE + SL) | 0.935 | +12.2% | 0.595 | 0.419 | 0.139 |
| (K + GE + SL) | 0.914 | +14.2% | 0.596 | 0.443 | 0.150 |
| (F) | 0.991 | +6.9% | 0.650 | 0.320 | 0.027 |
| (L) | 1.091 | -2.4% | 0.617 | 0.042 | -0.177 |
| (Ch) | 1.028 | +3.5% | 0.648 | 0.235 | 0.025 |
| (F + L) | 1.017 | +4.5% | 0.650 | 0.300 | 0.039 |
| (F + Ch) | 1.004 | +5.7% | 0.654 | 0.308 | 0.027 |
| (L + Ch) | 1.022 | +4.0% | 0.648 | 0.257 | 0.052 |
| (F + L + Ch) | 1.011 | +5.1% | 0.650 | 0.301 | 0.045 |
| Complete | 0.858 | +19.4% | 0.676 | 0.525 | 0.251 |

**Table 6:** a) G-*PLIP*’s performances with different graph topologies. ‘w/o PPI’ (*i*.*e*., protein-protein interaction) doesn’t take into account the protein-protein connectivity but uses the other available features and connections. ‘w/o CompSim’ (*i*.*e*., compound similarity) doesn’t use the compound-compound similarities but still integrates compound-compound connectivity (without weight on the edges). ‘w/o CompLink’ doesn’t connect compounds at all but use protein-protein connections. Similarly, ‘w/o PPI & CompLink’ uses neither protein-protein nor compound-compound connectivities. Each setup normally retains all other features and connections, and *−*Δ_*RMSE*_ is displayed compared to the complete model. ‘w/ DTIs’ (*i*.*e*., drug-target interaction) draws the drug-target interactions from the train set above every other connection mentioned, meaning the encoder won’t be limited to the protein or compound subgraph anymore. b) G-*PLIP*’s performances only using graph topologies. ‘only PPI’ utilizes only protein-protein connectivity. ‘only CompSim’ integrates compound-compound similarities. ‘only PPI & CompSim’ normally connects compounds and proteins. In all of these cases, other features are excluded, and *−*Δ_*RMSE*_ is reported in comparison to the null model.

| Model | $RMSE$ | $-\Delta_{RMSE}$ | $RBO$ | $R_p$ | $R^2$ |
| --- | --- | --- | --- | --- | --- |
| Complete | 0.858 | (*ref*) | 0.676 | 0.525 | 0.251 |
| w/o PPI | 0.857 | +0.1% | 0.686 | 0.534 | 0.262 |
| w/o CompSim | 0.883 | -2.9% | 0.677 | 0.509 | 0.209 |
| w/o CompLink | 0.870 | -1.4% | 0.679 | 0.521 | 0.223 |
| w/o PPI & CompLink | 0.883 | -2.9% | 0.685 | 0.517 | 0.212 |
| w/ DTIs | 0.854 | +0.5% | 0.693 | 0.541 | 0.269 |

(b) Graph informativeness, from the null model
| Model | $RMSE$ | $-\Delta_{RMSE}$ | $RBO$ | $R_p$ | $R^2$ |
| --- | --- | --- | --- | --- | --- |
| Null | 1.065 | (*ref*) | 0.596 | 0.061 | -0.034 |
| only PPI | 1.066 | -0.1% | 0.596 | 0.061 | -0.042 |
| only CompSim | 1.065 | 0.0% | 0.596 | 0.061 | -0.034 |
| only PPI & CompSim | 1.065 | 0.0% | 0.596 | 0.061 | -0.034 |

### 3.4 Informativeness of graph features

We assessed the performance of a series of partial versions of the model using combinatorial subsets of features. We started from the null model, a configuration encompassing protein-protein interactions and drug-drug similarities but without any specific node features. We then gradually added sets of protein or compound features to the null model. In each configuration, we documented the optimal performance achieved, thereby facilitating an exploration of the informativeness of individual features and their contributions to enhance predictive capacity.

According to the results (Table 5), *k*mer counts is the most pivotal feature, yielding a substantial improvement of RMSE by 14.08% compared to the null model. Other two protein features, gene expression and sub-cellular localization, do not noticeably contribute to the prediction beyond the capability of *k*mers.

We observed that protein node features emerge as more informative than their compound counterparts. As we suspected, molecular fingerprints improved the prediction. We also anticipated the observation that neither the features derived from Lipinski rules, nor other chemical features commonly used to roughly gauge physico-chemical properties of the molecules, exhibited prominent informativeness since they reflect rather on the pharmacokinetic properties and drug-likeliness of the molecules rather than bioactivity.

Considering that *k*mer was the most prominent feature that we identified, we attempted to extend or substitute *k*mer counts with ESM2 [81], a state-of-the-art embedding of proteins derived from evolutionary modelling. Yet we observed no discernible performance improvements S2, and the simplicity coupled with the good performance recommend *k*mer as an informative feature for any PLI prediction algorithm.

Another intriguing finding is that the model shows remarkably good performance even in the absence of protein-protein or compound-compound connectivity (Table 6). It’s essential to clarify that in such models, despite the absence of connections, the graph structure is used implicitly. This is because the encoder remains a key component of the model, and it is responsible for embedding the initial set of node features into a lower dimensional space. In addition, leaving out protein-protein and compound-compound connectivity leads to a small reduction in the performance. Nevertheless, we consider the observation interesting and worth reporting, because it underlies the power of GNNs to dissect relative contributions from node and edge features. Our attempt to model the protein-ligand interactions from the train set directly in the graph topology was not fruitful, and it suggests the presence of a restricted portion of the possible interactions is biasing the model.

## 4. Discussion

### 4.1 Learning capabilities of the model

Our structure-free approach compares favourably with advanced cutting-edge structure-aware models like MPNN-PLI, Pafnucy, K_*DEEP*_ or Fusion, as well as with novel structure-free models such as Yuel, when assessed in terms of RMSE.

Our evaluation highlights the importance of choosing an appropriate metric for benchmarking studies for the task of PLI prediction. As previously mentioned, the Pearson correlation coefficient can be misleading in evaluating regressive models, and we established that satisfactory scores can be achieved even by models incapable of predicting relevant values (Supplementary Figure S4).

GNNs offer a new approach to structure-free prediction of protein-ligand interaction. In docking-based approaches, the generation of poses is often one of the biggest bottlenecks in structural approaches, as it is challenging to generate plausible binding conformations of protein-ligand complexes without any experimental data. Our observations are consistent with recent studies discussing consistent conformational changes during drug-binding process [82], a challenge that current structural models are still striving to address.

The knowledge graph structure inherent to our methodology allows the encapsulation of informative protein and compound embeddings, obviating the need for an excessively complex model architecture. In this work, we only included a basic set of features in order to achieve a minimal and parsimonious model structure. Further work can build upon on our model to include more information thanks to the open and adaptive framework provided by GNNs.

The minimalistic architecture of G-*PLIP*, with only 8,449 trainable parameters, facilitates training on non-professional computing units without any high-end GPU and is robust to overfitting. In comparison, Yuel [76], the sole structure-free method tackling the same task we are aware of, comprises 1,411,523 trainable parameters.

### 4.2 Inference of the model

G-*PLIP* exhibits remarkable learning capabilities even in the absence of protein-protein or compound-compound connectivity, with the encoder layers being used to encapsulate the original features within a low-dimensional space without any convolutional mechanisms or aggregation. Regardless of the PPI database employed, and whether the PPI depicts physically interacting pairs only or a mix of functional and physical interactions, the protein-protein connectivity seems to make a limited contribution to the prediction. These observations, coupled with measures of node feature informativeness, suggest that the information aggregated from neighbouring nodes is currently not being effectively utilized by the model. One may wonder whether alternative designs of convolutional layers may better aggregate information from a biological knowledge graph. After all, previous reports and studies indicate that deep learning neural networks can encounter challenges when attempting to harness the learning potential of complex patterns they possess [51]. A further explanation could be found in the inherent shortcomings of the existing protein-protein interaction networks, particularly in their high rates of false positives and false negatives [83], and different PPI databases could be probed in this regard [84, 85]. We think that the protein-protein interaction is only one facet within a broader spectrum of relationships that a graph can depict. Other elements such as structural similarities, folding resemblance, or evolutionary distances could be explored with the architecture that we proposed.

Our GNN approach demonstrates its suitability for the task compared to standard machine learning models regarding to RMSE (Supp Table S1c). We therefore establish the graph architecture as an expressive framework capable of embedding a wide array of biological information. In other words, our work demonstrates that GNN is empowered to identify parsimonious models for a prediction task at hand. In line with this, we found that a restricted model using only *k*Mer and molecular fingerprints exhibits performances only slightly behind, 4.7% worse than the complete model. This also shown that our implemented features motivated by drug discovery purposes, namely gene expression (regardless of the tissue choice) and sub-cellular localization, offered a negligible contribution to the predictions. However, those results are meant to be investigated further, as we did not conduct complete experiments able to completely characterise the impact of individual features. While the trials with ESM2 did not yield improvements, standard protein and compound representations could still potentially be substituted by more advanced representation models (Table S2), for instance those leveraged by algorithms aiming at molecule properties prediction [86] or by protein language models tailored to capture specific characteristics such as coevolution [87, 88] or multiple sequence alignment-free embeddings [89].

Our observation of limited value of including lower-quality data confirms observations in previous studies [19, 79, 80]. Integrating data from biased or untidy biochemical assays carries a risk, even when performances on the dataset split initially appear promising. This underscores the necessity of systematic cross-validation against an external reference test set (in our case, the CASF 2016 Core set, remaining constant over the experiments). Evaluations on traditional random or temporal split can extend the biases of the original set.

### 4.3 Limitations

Biologically, G-*PLIP* performed comparably well across molecular function classes (Figure 3). The findings partially reflect the historical focus and current practice of small-molecule drug discovery. The relatively better performance for kinases, ion channels, and G protein-coupled receptors (GPCRs), may derive from the facts that many small molecules are discovered with such proteins as targets [90], and that they also comprise a large body of secondary pharmacological targets [91]. On the other side, some transcription factors (DNA-binding) are known to be challenging targets for small molecules [92]. The relatively variable performance of our model for multiple drug classes such as transcription factors, human hydrolases, and human proteases calls for alternative modelling strategies.

Methodologically, despite improved performances compared with recent models, G-*PLIP* occasionally demonstrates a propensity to underestimate potent bioactivities (Figure 2a), which remains critical for the identification of putative interactions to be explored by drug discovery projects. Besides, future work may explore ways to further improve G-*PLIP*’s performance, for instance performing pre-training on different properties (e.g. solubility measures).

Despite our efforts to investigate relative contributions of feature types, the interpretability of G-*PLIP* remains limited. Further analysis can be conducted to exploit the capability and limitations of the model. We may adopt a biology-informed split strategy, for instance splitting target proteins with hierarchical clustering, using pairwise structural and sequence homology of the proteins as distance metrics. We could also examine activity cliffs and their impacts on prediction, or scrutinize protein-ligand complexes leading to outlier predictions in order to devise improvement strategies.

Computationally, modelling PLI with a knowledge graph bears a significant memory cost. It may be mitigated by refactoring the data structure or by fine-tuning the compound similarity edges. These adjustments may improve memory efficiency and scalability.

## 5 Conclusion

Predicting protein-ligand interactions quantitatively represents an unsolved challenge at the core of small-molecule drug discovery. Here we present a structure-free approach that exhibits enhanced predictive accuracy compared to existing methods. Further, we demonstrate that information from the biochemical space can be encapsulated and leveraged by a deep learning procedure. Our methodology only requires an amino-acid sequence of proteins and SMILES representation of ligands, which makes it easily applicable. Despite its simplicity, G-*PLIP* exhibits remarkable learning and predictive capabilities.We demonstrate that G-*PLIP* can efficiently be trained from a relatively small qualitative dataset and yet proficiently extrapolate on unseen interactions from learned parameters. Composed of only a few layers, the model is swiftly trainable on non-specialized platforms and its computational performances position it as a suitable candidate for large-scale retraining or deployment.

## 6 Code availability and resource requirement

A reference implementation of G-*PLIP* platform, based on PyTorch [93] and PyTorch Geometric [94], is available at https://github.com/simoncrouzet/g-plip.

G-*PLIP* was trained with 16 cores of a Xeon-Gold CPU running at 2.1 GHz, with at least 64GB available RAM. It takes approximately 2,670 minutes to train G-*PLIP* on the PDBBind *complete* dataset, leading to a carbon footprint of 1.84 kg eCO2.

## Acknowledgements

This work has been conducted under the sponsorship of the internship program “Think Tank in Innovation and Sustainability #IP2TIS” initiated by André Hoffman, vice-chairman of the board of Roche Holding Ltd, and organized by Volker Herdtweck and colleagues, as well as the sponsorship of the Roche Advanced Analytics Network (RAAN). We thank Leo Klarner for esteemed discussions, alongside Nicolas Frey, Torsten Schindler, Francesco Brizzi and Christian Kramer for their valuable feedback, and colleagues from the Predictive Modeling and Data Analytics (PMDA) chapter of Pharmaceutical Sciences for their continuous support.

## Author contributions

S.J.C. conceived the idea, wrote the code, performed the experiments and wrote the manuscript. A.M.L. helped with the code writing and performed some experiments. K.A. wrote the manuscript and helped the project guidance with insightful discussions. L.S-P. and A.T.M. helped with the project guidance and the preparation of the manuscript with insightful discussions. T.N. contributed to the data gathering and helped to review the manuscript. M.DP. participated in reviewing the manuscript. J.D.Z. originated the idea, guided the project and wrote the manuscript.

