## Supplementary Material for "G-*PLIP*: Knowledge graph neural network for structure-free protein-ligand bioactivity prediction"

### 8 Supplementary Information

Table S1: a) Comparison of G-*PLIP*'s performances between several different PPI databases. b) Comparison of G-*PLIP*'s performances between global gene expressions and gene expression of a specific tissue (e.g. lung cells). c) Comparison of the architecture's performances of G-*PLIP* against standard machine learning algorithms, from the same (node) input features.

(a) PPI Databases

| **PPI Databases** | *RMSE* | *RBO* | $R_p$ | $R^2$ |
|---|---|---|---|---|
| Functional PPIs | 0.858 | 0.676 | 0.525 | 0.251 |
| Physical PPIs | 0.858 | 0.686 | 0.533 | 0.265 |
| Physical & Functional PPIs | 0.855 | 0.681 | 0.533 | 0.263 |

(b) Gene expression sets

| **Gene Expression Sets** | *RMSE* | *RBO* | $R_p$ | $R^2$ |
|---|---|---|---|---|
| Lung | 0.858 | 0.676 | 0.525 | 0.251 |
| All Genes | 0.867 | 0.674 | 0.526 | 0.244 |

(c) Stand-off against state-of-the-art machine learning techniques from AutoGluon, a package providing quick deployment and optimization of a high-accuracy machine learning model. Over an ensemble of learning algorithms, AutoGluon identifies 'WeightedEnsemble_L2' as the top-performing one.

| **Algorithms** | *RMSE* | $R_p$ | $R^2$ |
|---|---|---|---|
| G-*PLIP* | 0.862 | 0.525 | 0.246 |
| AutoGluon | 1.015 | 0.631 | 0.398 |

Table S2: Informativeness of sequence encodings, comparing the informativeness of $k$Mer and ESM2 embeddings.

| **Sequence encoding** | *RMSE* | *RBO* | $R_p$ | $R^2$ |
|---|---|---|---|---|
| *no Encoding* | 0.915 | 0.663 | 0.458 | 0.183 |
| $k$Mer | 0.859 | 0.696 | 0.525 | 0.244 |
| ESM2 | 0.868 | 0.675 | 0.524 | 0.253 |
| $k$Mer + ESM2 | 0.863 | 0.676 | 0.526 | 0.262 |

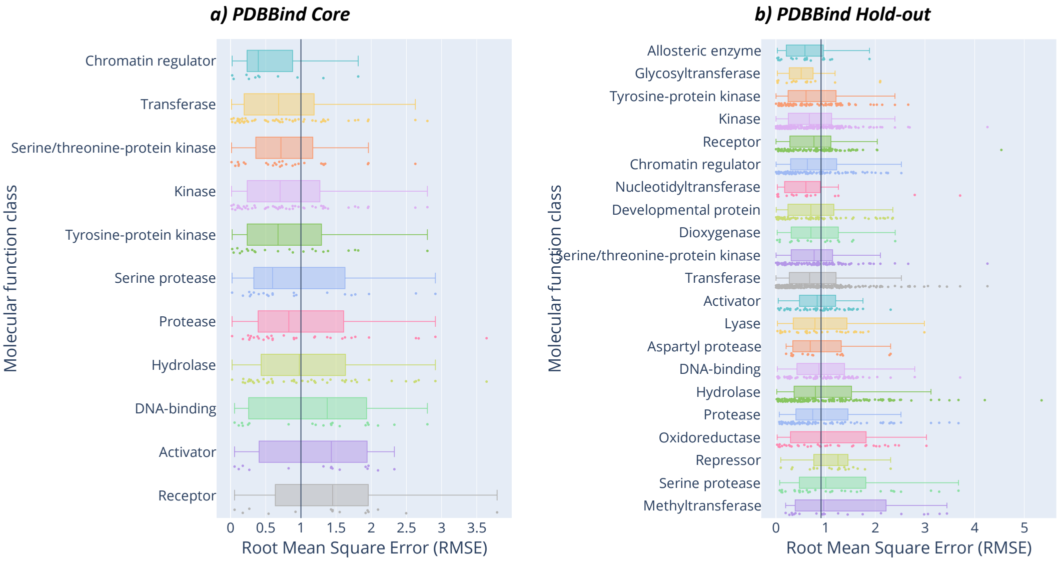


Figure S1: Model's performance on the PDBBind Core test set (panel a) and PDBBind Hold-out test set (panel b) across different molecular function classes of targets. Target classes are sorted by median $RMSE$ values. The black vertical line is the overall mean value across all molecular function classes. Molecular functions possessing less than 10 targets (for the PDBBind Core test set) or 16 targets (for the PDBBind Hold-out test set) are not displayed.

Table S3: Parameters benchmarked during G-*PLIP*'s design process.

| **Parameters** | **Values** |
|---|---|
| Number of convolutional layers | {1, 2, 3, 4} |
| Type of convolutional layers | {SAGE, GAT} |
| Learning rate | {0.01, 0.005, 0.001, 0.0005, 0.0001} |
| Batch size | {256, 512, 1024} |
| Dropout | {0.0, 0.1, 0.2}; {$in\_encoder$, $in\_decoder$, $in\_both$} |
| Compound similarity threshold | {0.0, 0.5, 0.8} |
| Morgan fingerprints size | {128, 256, 512, 1024} |

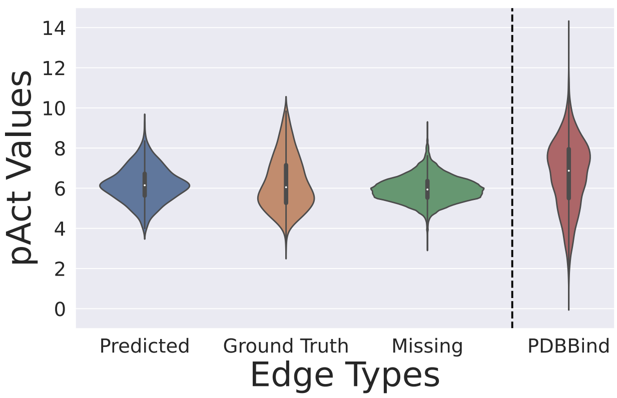


Figure S2: Distributions of *pAct* values over diverse setups of protein-ligand interactions. From left to right: Predictions of labelled edges, Ground truths of labelled edges, and predictions of unlabelled edges. Unlabelled edges do not necessarily mean that the protein-ligand pair does not interact; it may also mean that the interaction has not been tested or reported.

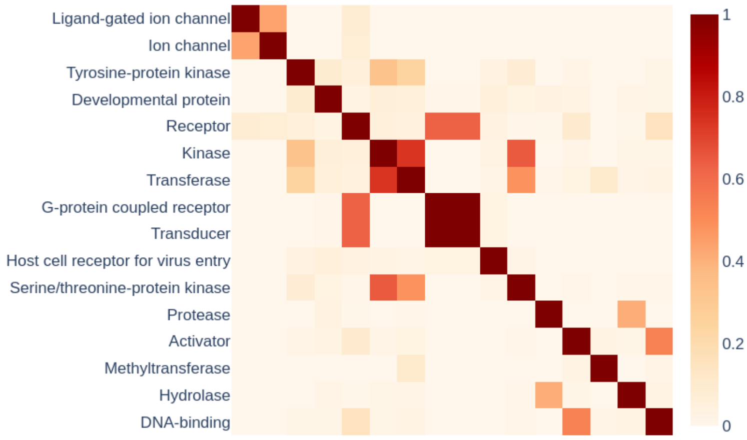


Figure S3: Displayed heatmap shows the Jaccard similarity index ranging from 0 to 1. The higher the Jaccard index the more similar are the two molecular function classes in terms of shared targets.

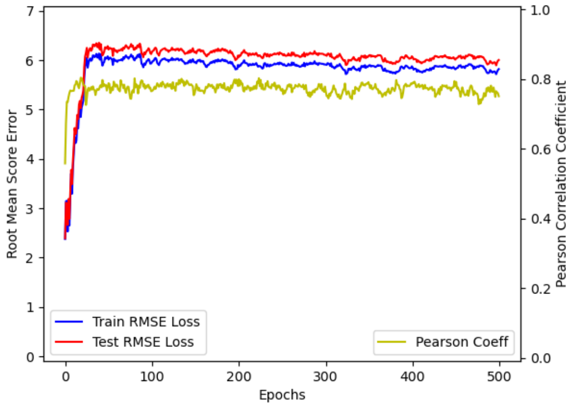


Figure S4: Learning curve of a training aiming to optimize the Pearson correlation coefficient. Despite a correct score, the model totally fails to grasp any useful knowledge from the data, as shown by the RMSE values.
